# Distinct Glutamatergic Inputs to the Nucleus Accumbens Differentially Regulate Vulnerability to Cannabinoid Addiction

**DOI:** 10.64898/2026.07.17.739252

**Authors:** María Fernanda Ponce-Beti, Tatiana Gusinskaia, Ignacio Marin-Blasco, Roberto Capellán, Mariana G. Fronza, Raul Andero, Rafael Maldonado, Elena Martín-García

**Affiliations:** Laboratory of Neuropharmacology-Neurophar, Department of Medicine and Life Sciences, Universitat Pompeu Fabra (UPF), Barcelona, Spain; Institut de Neurociències, Universitat Autònoma de Barcelona, Cerdanyola del Vallès, Barcelona, Spain; Departament de Psicobiologia i de Metodologia de les Ciències de la Salut. Universitat Autònoma de Barcelona, Cerdanyola del Vallès, Barcelona, Spain; Centro de Investigación Biomédica En Red en Salud Mental (CIBERSAM), Instituto de Salud Carlos III, Madrid, Spain; Unitat de Neurociència Traslacional, Parc Taulí Hospital Universitari, Institut d’Investigació i Innovació Parc Taulí (I3PT), Institut de Neurociències, Universitat Autònoma de Barcelona, Cerdanyola del Vallès, Spain; ICREA, Barcelona, Spain; Fundació Institut Hospital del Mar d’Investigacions Mèdiques (HMRIB), Barcelona, Spain

**Author notes:** These authors contributed equally to this work (María Fernanda Ponce-Beti, Tatiana Gusinskaia). These authors equally supervised this work (Rafael Maldonado, Elena Martín-García).

## Abstract

Cannabis use disorder (CUD) is a chronic relapsing disorder characterized by compulsive drug seeking, persistent drug use despite adverse consequences, and a high risk of relapse. Although the nucleus accumbens (NAc) is a central hub in the neural circuitry underlying addiction, the specific glutamatergic inputs regulating vulnerability to cannabinoid addiction remain poorly understood. Here, we investigated the contribution of two major limbic glutamatergic projections to the NAc, the dorsal hippocampus (dHPC) to NAc and basolateral amygdala (BLA) to NAc pathways, using a validated mouse model of WIN55,212-2 intravenous self-administration combined with projection-specific chemogenetic inhibition. Male C57BL/6J mice received combinatorial viral vector delivery of inhibitory hM4Di DREADDs selectively targeting either the dHPC to NAc or the BLA to NAc pathway. Chronic pathway inhibition was achieved by continuous administration of deschloroclozapine through osmotic minipumps during the development of cannabinoid addiction-like behavior. Animals were evaluated using a multidimensional behavioral paradigm assessing the three-core addiction-like criteria of persistence of drug seeking, motivation, and compulsive-like behavior, as well as craving-related behaviors and phenotypic vulnerability traits. Chronic inhibition of either the dHPC to NAc or the BLA to NAc pathway significantly increased the proportion of mice developing an addiction-like phenotype. Both manipulations enhanced persistence of drug seeking during periods of drug unavailability, identifying persistence as a shared behavioral consequence of disrupting glutamatergic signaling to the NAc. In contrast, the two pathways differentially regulated other addiction-related behaviors. Inhibition of the dHPC to NAc pathway increased motivation to obtain WIN55,212-2, impulsivity, reward sensitivity, and resistance to extinction, whereas inhibition of the BLA to NAc pathway selectively enhanced cue-induced drug seeking. Neither manipulation altered compulsive-like responding, locomotor activity, or body weight. These findings demonstrate that distinct glutamatergic afferents to the NAc differentially regulate vulnerability to cannabinoid addiction-like behavior while converging on persistence as a common circuit-level mechanism. Our results establish the NAc as an integrative hub coordinating complementary contextual and emotional information during the transition to cannabinoid addiction and provide a circuit-based framework for understanding the neural mechanisms underlying Cannabis Use Disorder.

## 1. INTRODUCTION

Cannabis is the most widely used illicit psychoactive substance worldwide, and its consumption continues to increase as social acceptance and legalization expand across many countries (Freeman et al., 2026). Recent epidemiological reports indicate that approximately 8–9% of European adults consumed cannabis during the previous year, with substantially higher rates among adolescents and young adults (European Monitoring Centre for Drugs and Drug Addiction, 2026). Although cannabis use is common, only a subset of users develops Cannabis Use Disorder (CUD), a severe psychiatric condition characterized by compulsive drug seeking and drug use despite adverse consequences (Hall et al., 2026). Understanding why some individuals remain resilient, whereas others become vulnerable to addiction, represents one of the major challenges in addiction neuroscience.

The transition from recreational drug use to addiction is not a linear process but rather a progressive selection of vulnerable individuals along the trajectory from initial drug exposure to compulsive drug use (Piazza and Deroche-Gamonet, 2013). Among those who initiate cannabis consumption, only a proportion progresses from occasional use to regular use, and an even smaller proportion ultimately develops CUD. This transition to addiction results from a complex interplay among genetic susceptibility, environmental influences, and epigenetic mechanisms that shape brain function. According to the Diagnostic and Statistical Manual of Mental Disorders, Fifth Edition (DSM-5), CUD is diagnosed based on eleven clinical criteria that reflect impaired control over drug use, compulsive drug seeking, and persistent consumption despite harmful consequences (American Psychiatric Association, 2022). These clinical features can be modeled experimentally using multidimensional animal paradigms that evaluate three core addiction-like behaviors: persistence of drug seeking, increased motivation for the drug, and compulsive-like responding despite punishment (Maldonado et al., 2021).

Current neurobiological models conceptualize addiction as a cyclic disorder involving three interconnected stages: binge/intoxication, withdrawal/negative affect, and preoccupation/anticipation (Koob and Volkow, 2016). These behavioral stages are supported by partially overlapping neural circuits involving the basal ganglia, the extended amygdala, and the prefrontal cortex, which undergo progressive neuroadaptations during repeated drug exposure (Hoch et al., 2025). Within this framework, the nucleus accumbens (NAc) occupies a central position by integrating glutamatergic information from multiple cortical and limbic regions to regulate reward processing, reinforcement learning, motivation, and drug-seeking behavior (Volkow et al., 2019). Although considerable attention has focused on corticostriatal circuits, considerably less is known about how distinct glutamatergic afferents to the NAc contribute to individual vulnerability to cannabinoid addiction.

Previous work from our laboratory demonstrated that glutamatergic signaling within the prelimbic cortex (PL) to NAc pathway critically regulates vulnerability to food addiction (Domingo-Rodriguez et al., 2020). Genetic deletion of cannabinoid CB1 receptors from cortical glutamatergic neurons promoted resilience to addiction-like behavior, whereas chemogenetic inhibition of the PL to NAc pathway or overexpression of dopamine D2 receptors within this circuit shifted animals toward a vulnerable phenotype characterized by enhanced compulsive-like behavior. These findings established the NAc as a key hub controlling the transition from resilience to vulnerability, and raised the possibility that other glutamatergic inputs converging onto the NAc regulate distinct components of addiction.

Among the major excitatory afferents to the NAc, the dorsal hippocampus (dHPC) and the basolateral amygdala (BLA) are particularly relevant (Volkow et al., 2019). The dHPC provides contextual and spatial information that contributes to reinforcement learning, contextual memory, and habit formation, processes primarily associated with the binge/intoxication stage of addiction (Koob and Volkow, 2016). In contrast, the BLA conveys emotional and motivational information involved in reward valuation, cue-associated learning, and relapse, and has been strongly implicated in the negative affect and preoccupation/anticipation stages of the addiction cycle. Despite their established anatomical connectivity with the NAc, the specific contribution of these glutamatergic pathways to the development of cannabinoid addiction remains unknown.

In the present study, we tested the hypothesis that the dHPC-to-NAc and BLA-to-NAc glutamatergic projections differentially regulate vulnerability to cannabinoid addiction. Using a projection-specific chemogenetic manipulation strategy combined with intravenous WIN55,212-2 self-administration, a validated preclinical model of cannabinoid addiction-like behavior (Cajiao-Manrique et al., 2023b), we selectively inhibited each pathway throughout the development of addiction-like behavior. We evaluated the impact of each manipulation on the three addiction-like criteria, craving-related behaviors, and phenotypic vulnerability traits. By dissecting the contribution of these major limbic glutamatergic inputs to the NAc, this study provides new mechanistic insights into the circuit-level regulation of vulnerability to cannabinoid addiction and identifies these pathways as potential targets for future therapeutic interventions.

## 2. EXPERIMENTAL PROCEDURES

### 2.1. Animals

39 eight-week-old male C57BL/6J mice (Charles River Laboratories, France) were housed individually under controlled environmental conditions (21 ± 1°C; 55 ± 10% humidity) with food and water available *ad libitum*. Animals were maintained under a reversed 12-h light/dark cycle (lights off at 08:00 h), and all behavioral experiments were conducted during the first hours of the dark phase. Body weight and food intake were monitored throughout the study. All experimental procedures were approved by the local Animal Ethics Committee (Comitè Ètic d’Experimentació Animal–Parc de Recerca Biomèdica de Barcelona, CEEA-PRBB, CEEA 1667-RML-22-0076-P1) and conducted in accordance with the European Communities Council Directive 2010/63/EU regulating animal experimentation. Mice were randomly assigned to one of the three experimental groups: (1) Control (n = 12), (2) dHPC to NAc (n = 16), or (3) BLA to NAc (n = 11).

### 2.2. Drugs

The synthetic cannabinoid receptor full agonist WIN55,212-2 mesylate salt (Sigma-Aldrich, St. Louis, MO, USA) was dissolved in one drop of Tween-80 (50 μL) and diluted in sterile physiological saline. Mice received a single intraperitoneal injection of WIN55,212-2 at a dose of 0.1 mg/kg, 24 h before the first operant self-administration session to prevent the aversive effects associated with the initial cannabinoid exposure (Mendizábal et al., 2006). During self-administration, WIN55,212-2 was delivered intravenously at a dose of 12.5 μg/kg per infusion in a volume of 23.5 μL over 2 s (Vallee et al., 2014). All solutions were strictly protected from light and stored at room temperature throughout the duration of the experimental procedures.

Deschloroclozapine (DCZ) was continuously administered using subcutaneous osmotic minipumps (Alzet Model 2004, DURECT Corporation, Cupertino, CA, USA). Osmotic minipumps were filled with either vehicle or DCZ at a final concentration of 0.5 mg/mL in sterile physiological saline and implanted during jugular vein catheterization surgery, providing continuous drug delivery at a dose of 0.1 mg/kg/day throughout the 28-day behavioral protocol.

### 2.3. Viral vectors

Projection-specific chemogenetic inhibition of glutamatergic inputs to the NAc was achieved using a dual-viral strategy combining a retrogradely transported Cre recombinase-expressing AAV —AAVretro-pmSyn1-EBFP-Cre (8.2E+12 gc/ml)— with a Cre-dependent inhibitory DREADD-expressing AAV —AAV8-CamKII-DIO-hM4Di(Gi)-mCherry (1.21E+13 gc/ml)—, both produced at the Viral Vector Production Unit of the Universitat Autònoma de Barcelona (UPV-UAB). Control mice received the corresponding Cre-dependent vector —AAV8-hSyn-DIO-mCherry (1.4E+13 gc/ml) —.

### 2.4 Stereotaxic surgery and chemogenetic manipulation of dHPC to NAc and BLA to NAc projections

Mice were anesthetized with ketamine hydrochloride (75 mg/kg, i.p.) and medetomidine hydrochloride (1 mg/kg, i.p.) dissolved in sterile physiological saline and positioned in a stereotaxic apparatus (Kopf Instruments, USA). Bilateral viral injections were performed using 33-gauge stainless-steel injectors connected through PE-20 polyethylene tubing to a 10-μL Hamilton microsyringe.

For the dHPC to NAc group, AAV8-CamKII-DIO-hM4Di(Gi)-mCherry was bilaterally injected into the dHPC, whereas AAVretro-pmSyn1-EBFP-Cre was bilaterally injected into the NAc core. For the BLA to NAc group, the same Cre-dependent hM4Di vector was bilaterally injected into the BLA together with AAVretro-pmSyn1-EBFP-Cre injections into the NAc core. Control mice received identical stereotaxic injections but using the AAV8-hSyn-DIO-mCherry as Cre-dependent vector (Domingo-Rodriguez et al., 2020). The Cre-dependent viral vector was infused at a rate of 0.05 μL/min (0.20 μL/site), whereas the retrograde vector was delivered at 0.10 μL/min (0.40 μL/site). After each infusion, the injector remained in place for 10 min to facilitate viral diffusion and minimize reflux before being slowly withdrawn. Stereotaxic coordinates were determined according to the Paxinos and Franklin mouse brain atlas (Paxinos, 2019). Viral injections targeted the dHPC, BLA, and NAc core subdivision using the following coordinates: dHPC: AP −2.00 mm, ML ±1.50 mm, DV −1.45 mm; BLA: AP −1.34 mm, ML ± 2.9 mm, DV −4.80 mm; NAc core: AP +1.94 mm, ML ±1.00 mm, DV −4.60 mm. Body temperature was maintained at 36°C using a thermostatically controlled heating pad throughout the surgical procedures. Animals remained undisturbed during four weeks to allow recovery and stable viral expression before behavioral testing.

### 2.5. Jugular vein catheterization

Following viral surgery and the recovery period, mice were implanted with chronic indwelling intravenous catheters in the right jugular vein as previously described (Martín-García et al., 2026). Briefly, mice were anesthetized with ketamine (75 mg/kg, i.p.) and medetomidine (1 mg/kg, i.p.), and a silastic catheter connected to a stainless-steel cannula mounted on a dental cement pedestal was inserted into the right jugular vein. The external cannula exited at the mid-scapular region and allowed intravenous drug delivery during operant sessions. Following surgery, mice received gentamicin (1 mg/kg, i.p.), meloxicam (2 mg/kg, s.c.), glucose saline solution, and atipamezole (2.5 mg/kg, s.c.) to reverse anesthesia. At the same surgical procedure, osmotic minipumps containing DCZ or vehicle were implanted subcutaneously in the lower back. Animals were allowed to recover for three days before the initiation of operant self-administration. Catheter patency was evaluated at the end of the behavioral protocol using intravenous administration of sodium thiopental (5 mg/mL). A functional catheter was confirmed by the rapid onset of ataxia and loss of muscle tone within 5–10 s after injection. Animals failing to meet this criterion were considered to have non-patent catheters and were excluded from the analyses. To maintain catheter patency, they were flushed daily with heparinized saline after each self-administration session.

### 2.6. Operant conditioning maintained by WIN55,212-2 intravenous self-administration

Operant self-administration was performed in standard mouse operant conditioning chambers (ENV-307A-CT, Med Associates, St. Albans, VT, USA) equipped with two nose-poke holes, one designated as active and the other as inactive (Martín-García et al., 2026). Responses in the active hole resulted in intravenous delivery of WIN55,212-2 paired with illumination of the cue light located above the active nose-poke and the sound of the infusion pump, whereas responses in the inactive hole had no programmed consequences. Twenty-four hours before the first operant session, mice received a single intraperitoneal injection of WIN55,212-2 (0.1 mg/kg) to minimize the aversive effects associated with the initial cannabinoid exposure. Animals were then trained to self-administer WIN55,212-2 (12.5 μg/kg/infusion) under a fixed-ratio 1 (FR1) schedule of reinforcement for five consecutive daily sessions followed by a fixed-ratio 2 (FR2) schedule for an additional five sessions. Each daily session lasted 125 min and consisted of two 55-min drug-available periods separated by a 15-min drug-free interval. At the beginning of each session, mice received one non-contingent priming infusion of WIN55,212-2. Following each infusion, a 10-s timeout period was imposed during which additional responses had no programmed consequences.

The session ended after 50 reinforcers had been obtained or after 125 min, whichever occurred first. Acquisition criteria were considered achieved when animals fulfilled the following conditions over three consecutive sessions: (i) less than 20% variability in the number of reinforcers obtained, (ii) at least 75% of total responses directed toward the active nose-poke, and (iii) a minimum of five reinforcers per session (Martín-García et al., 2026).

### 2.7. Assessment of addiction-like behaviors

Following the acquisition, mice were evaluated for three addiction-like criteria adapted from the multidimensional model of addiction-like behavior described by Deroche-Gamonet and colleagues (Deroche-Gamonet et al., 2004) and evaluated across the three core addiction-like behavioral hallmarks adapted from the DSM-5 substance use criteria (Cajiao-Manrique et al., 2023a; Cajiao-Manrique et al., 2023b; Martín-García et al., 2026). Animals were classified according to persistence of drug seeking, motivation for the drug, and compulsive-like responding. For each behavioral criterion, the threshold for a positive score was established as the 75th percentile of the distribution obtained in the control group (AAV8-hSyn-DIO-mCherry). Mice fulfilling two or three criteria were classified as addicted, whereas those fulfilling zero or one criterion were classified as non-addicted.

#### 2.7.1. Persistence of drug seeking

Persistence was evaluated as the number of responses performed in the active nose-poke during the 15-min drug-free period (DFP) separating the two drug-available periods of the self-administration session. This parameter was measured during the three consecutive sessions preceding the progressive ratio test, and the average value was used for statistical analyses.

#### 2.7.2. Motivation

Motivation to obtain WIN55,212-2 was assessed under a progressive-ratio (PR) schedule of reinforcement. The PR was conducted in the self-administration chambers on the day following the fifth FR2 session. The response requirement increased according to the following progression: 1, 5, 12, 21, 33, 51, 75, 90, 120, 155, 180, 225, 260, 300, 350, 410, 465, 540, 630, 730, 850, 1000, 1200, 1500, 1800, 2100, 2400, 2700, 3000, 3400, 3800, 4200, 4600, 5000, and 5500 responses. The breakpoint was defined as the highest completed ratio before the animal ceased responding. Sessions ended after 4 h or after 1 h without responding.

#### 2.7.3. Compulsive-like responding

Compulsivity was evaluated in a shock session in which drug seeking was associated with an aversive consequence. It was assessed following an additional FR2 session to re-establish baseline behavior after the PR test. Mice were randomly assigned to a different self-administration chamber while maintaining their original active-hole side to avoid context-specific habituation. Animals were tested under an FR2 schedule for 50 min. The first active response triggered a mild foot shock (0.18 mA, 2 s), whereas completion of the second response delivered both the foot shock and the WIN55,212-2 infusion together with the associated cue light. If the second response was not completed within 60 s after the first response, the sequence was reset. The total number of shocks received during the session was used as the measure of compulsive-like responding.

### 2.8. Parameters related to craving: Extinction and cue-induced reinstatement

Following completion of the addiction-like behavioral tests, mice underwent daily 2-h extinction sessions during which responses in the active nose-poke no longer resulted in drug delivery or presentation of the associated cues. Extinction continued until animals reached the extinction criterion, defined as fewer than 35% of the active responses performed during the final three self-administration sessions across three consecutive extinction sessions.

#### 2.8.1. Resistance to extinction

Resistance to extinction was quantified as the number of active nose-poke responses during the first extinction session.

#### 2.8.2. Cue-induced reinstatement

Following 20 extinction sessions, animals were exposed to a cue-induced reinstatement test. Two consecutive active nose-pokes triggered the cue light and the sound associated with drug delivery, although no infusion was administered. Drug-seeking was reported as the total number of active nose-pokes during the 2-hour session.

### 2.9. Phenotypic vulnerability traits

#### 2.9.1. Impulsivity

Impulsivity-like behavior was quantified as the number of active responses emitted during the 10-s timeout period following each WIN55,212-2 infusion. Values were averaged across the three self-administration sessions preceding the progressive-ratio test.

#### 2.9.2. Sensitivity to reward

Reward sensitivity was assessed as the average number of reinforcers obtained during the final three FR2 self-administration sessions.

### 2.10. Animal classification

Mice were classified into “addicted” (AD) and “non-addicted” (NA) groups based on the number of addiction-like criteria met (motivation, persistence, and compulsivity). An animal was considered positive for a criterion if its score was at or above the 75th percentile of the total population distribution. Mice meeting 0 or 1 criterion were classified as non-addicted, while those meeting 2 or 3 criteria were classified as addicted.

### 2.11. Histological verification

At the completion of the behavioral experiments, mice were deeply anesthetized and transcardially perfused with phosphate-buffered saline followed by 4% paraformaldehyde. Brains were post-fixed, cryoprotected, and sectioned coronally. Viral expression and injection sites were verified by fluorescence microscopy. Animals showing inaccurate viral expression or off-target injections were excluded from the final analyses.

### 2.12. Statistical analysis

Behavioral data were analyzed using SPSS Statistics (IBM) and GraphPad Prism (GraphPad Software). Normality was assessed using the Kolmogorov–Smirnov test. Data were analyzed using one-way or two-way ANOVA with repeated measures when appropriate, followed by Tukey’s or Fisher’s post hoc tests for multiple comparisons. Non-parametric data were analyzed using the Mann–Whitney U test or Kruskal–Wallis test followed by Dunn’s multiple-comparison test. Categorical variables were compared using χ^2^ tests. Pearson correlation analyses were used to evaluate associations between addiction-like behavioral measures. Results are expressed as mean ± SEM or median with interquartile range, as indicated in the figure legends. Statistical significance was set at *P* < 0.05.

## 3. RESULTS

### 3.1. Chemogenetic inhibition of dHPC to NAc and BLA to NAc projections increases vulnerability to cannabinoid addiction-like behavior

A projection-specific chemogenetic approach was used to selectively inhibit the dHPC to NAc or the BLA to NAc pathways by combining retrograde AAV-Cre delivery into the NAc with Cre-dependent hM4Di expression in CaMKII-expressing neurons of the dHPC or BLA during a WIN55,212-2 intravenous self-administration protocol (Fig. 1A,B). During FR1 and FR2 training, all groups acquired operant responding maintained by WIN55,212-2 self-administration, as shown by the progressive increase in active nose-poke responses across sessions. Notably, inhibition of the dHPC to NAc pathway showed a significantly higher number of active nose-pokes during training compared with the Control group (Fig. 2A; repeated-measures ANOVA followed by Dunnett’s post hoc test, p = 0.020).

**Figure 1.**
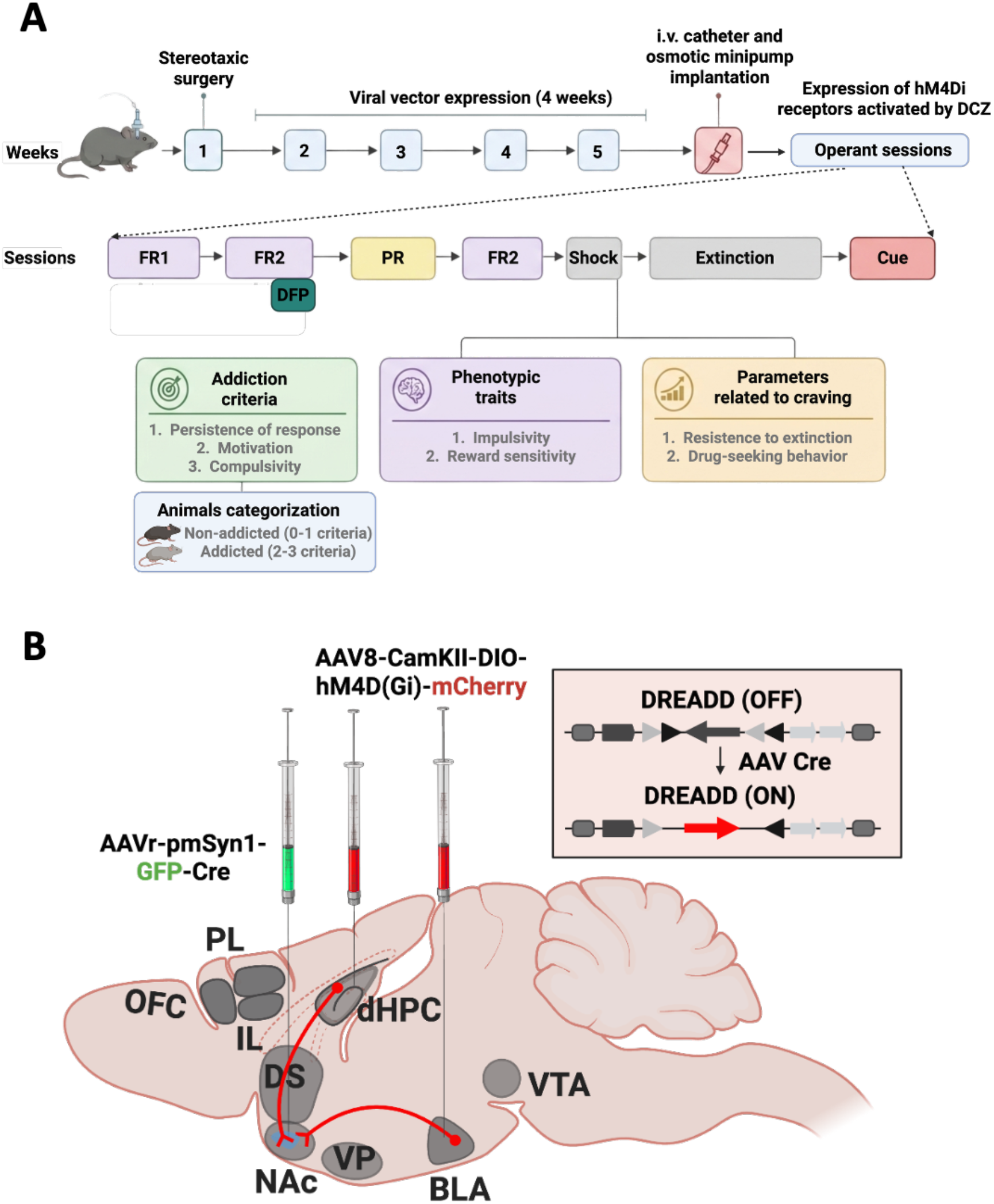
Experimental design and projection-specific chemogenetic inhibition of dHPC to NAc and BLA to NAc pathways. **(A)** Experimental timeline. Mice underwent stereotaxic surgery followed by four weeks of viral expression. Intravenous catheter implantation and osmotic minipump implantation were subsequently performed before the initiation of WIN55,212-2 intravenous self-administration. Behavioral testing consisted of five fixed-ratio 1 (FR1) sessions, five fixed-ratio 2 (FR2) sessions, a progressive-ratio (PR) test, one additional FR2 session, a shock-resistance test, extinction sessions, and a cue-induced reinstatement test. Addiction-like behavior was evaluated using three behavioral criteria (persistence of response, motivation, and compulsive-like behavior). Animals were classified as non-addicted (0–1 criteria) or addicted (2–3 criteria). Impulsivity and reward sensitivity were assessed as phenotypic vulnerability traits, whereas resistance to extinction and cue-induced reinstatement were evaluated as parameters related to craving. **(B)** Schematic representation of the combinatorial chemogenetic strategy. A retrogradely transported AAV expressing Cre recombinase (AAVretro-Syn-GFP-Cre) was injected into the nucleus accumbens (NAc), whereas a Cre-dependent inhibitory DREADD vector (AAV8-CamKII-DIO-hM4Di(Gi)-mCherry) was injected into either the dorsal hippocampus (dHPC) or the basolateral amygdala (BLA), thereby restricting hM4Di expression to dHPC or BLA glutamatergic neurons projecting to NAc.

**Figure 2.**
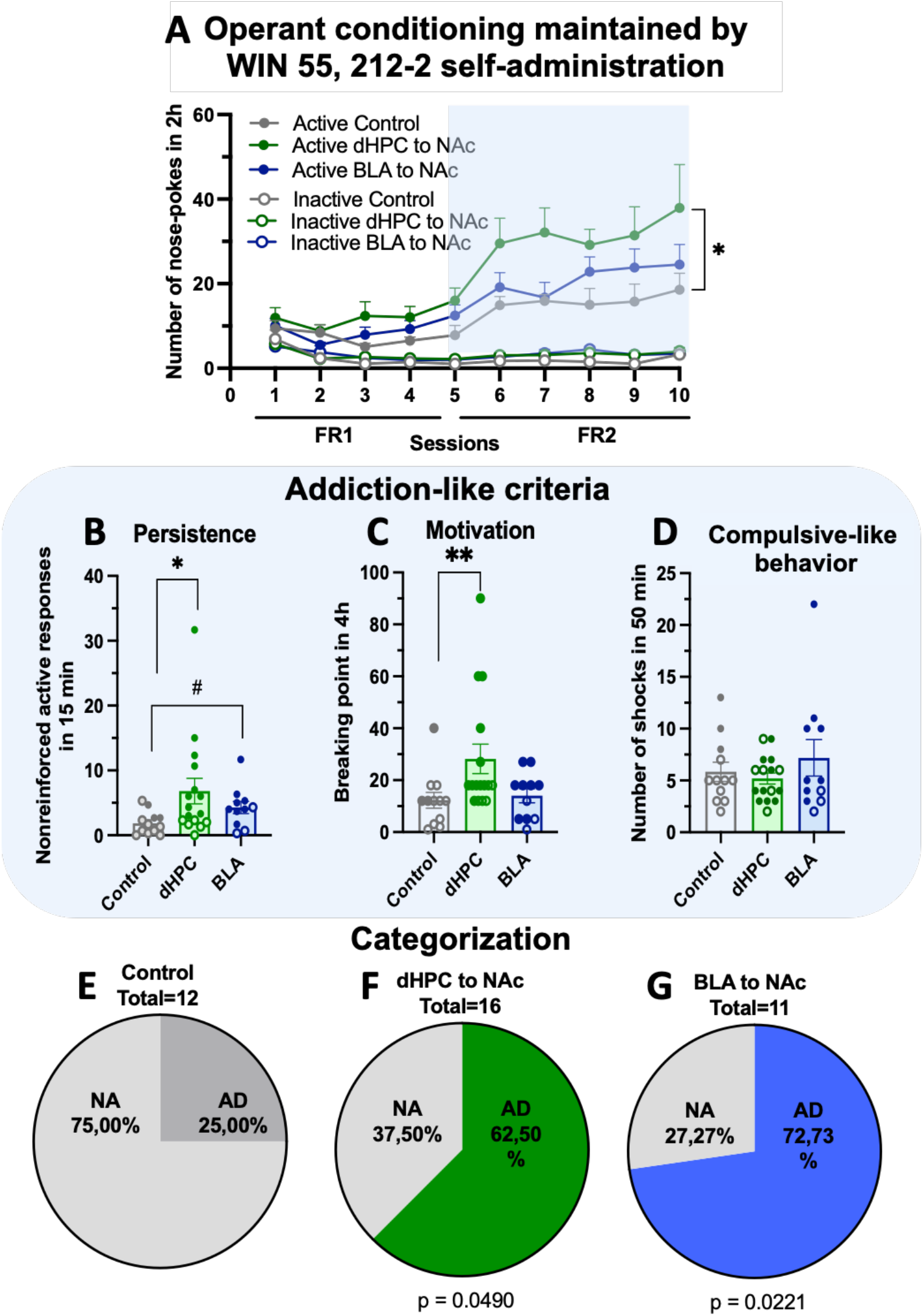
Chemogenetic inhibition of dHPC to NAc and BLA to NAc projections increases vulnerability to cannabinoid addiction-like behavior. **(A)** Active and inactive nose-poke responses during WIN55,212-2 self-administration under FR1 and FR2 schedules of reinforcement (two-way repeated-measures ANOVA). **(B–D)** Assessment of the three addiction-like behavioral criteria. **(B)** Persistence of response, measured as nonreinforced active responses during the 15-min drug-free period. **(C)** Motivation, measured as the breakpoint achieved during the progressive-ratio test. **(D)** Compulsive-like behavior, measured as the number of shocks received during the punishment session. Filled circles represent addicted mice and open circles represent non-addicted mice. **(E)** Classification of animals according to addiction-like behavior. Pie charts show the percentage of non-addicted (NA; 0–1 criteria) and addicted (AD; 2–3 criteria) mice in each experimental group. *p<0.05, **p<0.01 for statistical differences from repeated measures ANOVA, t-test, or Mann-Whitney U, depending on the normality test. Mice were randomly assigned to one of the three experimental groups: (1) Control (n = 12), (2) dHPC to NAc (n = 16), or (3) BLA to NAc (n = 11). Data are presented as mean ± SEM.

Assessment of the three addiction-like criteria revealed pathway-specific behavioral effects. Inhibition of the dHPC to NAc pathway significantly increased persistence of drug seeking during the drug-free period (Fig. 2B; Mann–Whitney U test, p = 0.013) and motivation for WIN55,212-2, as measured by the progressive-ratio breakpoint (Fig. 2C; Mann–Whitney U test, p = 0.005). Inhibition of the BLA to NAc pathway also increased persistence of drug seeking compared with the control group (Fig. 2B; t-test, p = 0.031). No significant differences between groups were observed in compulsive-like responding during the punishment test (Fig. 2D).

The proportion of mice classified as addicted was significantly increased in both pathway-inhibited groups compared with the Control group (Fig. 2E). Specifically, 25.00% of Control mice met the criteria for addiction (3/12), whereas this proportion increased to 62.50% in the dHPC to NAc group (10/16; χ^2^ test, p = 0.049) and to 72.73% in the BLA to NAc group (8/11; χ^2^ test, p = 0.022).

Together, these results indicate that chronic inhibition of glutamatergic dHPC to NAc and BLA to NAc projections enhances vulnerability to cannabinoid addiction-like behavior, with partially distinct effects on individual addiction-like criteria.

### 3.2. dHPC to NAc and BLA to NAc inhibition differentially affects craving-related parameters and phenotypic vulnerability traits

We next evaluated whether inhibition of dHPC to NAc and BLA to NAc projections altered behavioral dimensions related to craving and addiction vulnerability. Mice in the dHPC to NAc group showed significantly increased impulsivity-like behavior, measured as active responses during the timeout period (Fig. 3B; Mann–Whitney U test, p = 0.026), and increased sensitivity to reward, measured as the number of reinforcers obtained during the final self-administration sessions (Fig. 3C; Mann–Whitney U test, p = 0.002). Regarding craving-related parameters, inhibition of the dHPC to NAc pathway significantly increased resistance to extinction, measured as active responses during the first extinction session (Fig. 3D; t-test, p = 0.031). In contrast, inhibition of the BLA to NAc pathway selectively increased cue-induced drug-seeking behavior during reinstatement (Fig. 3E; Mann–Whitney U test, p = 0.037).

**Figure 3.**
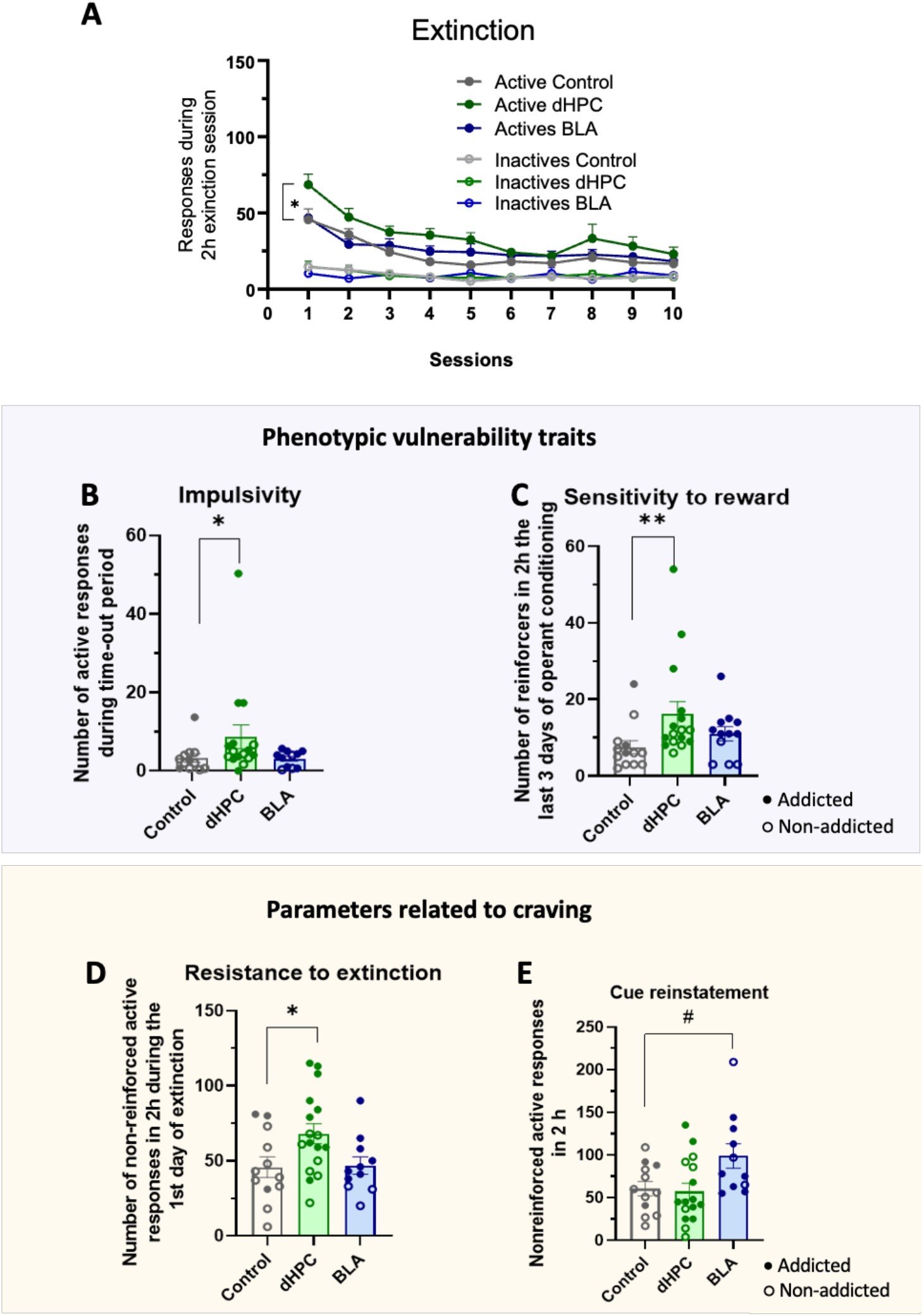
Chemogenetic inhibition of dHPC to NAc and BLA to NAc projections differentially regulates craving-related behaviors and phenotypic vulnerability traits. **(A)** Active and inactive responses during extinction sessions. **(B–C)** Phenotypic vulnerability traits. **(B)** Impulsivity, measured as active responses during the timeout period. **(C)** Reward sensitivity, measured as the number of reinforcers obtained during the final three FR2 sessions. **(D–E)** Craving-related parameters. **(D)** Resistance to extinction, measured as active responses during the first extinction session. **(E)** Cue-induced reinstatement, measured as nonreinforced active responses during the reinstatement test. Filled circles represent addicted mice and open circles represent non-addicted mice. *p<0.05, **p<0.01, #p<0.05 for statistical differences from repeated measures ANOVA, t-test, or Mann-Whitney U, depending on the normality test. Mice were randomly assigned to one of the three experimental groups: (1) Control (n = 12), (2) dHPC to NAc (n = 16), or (3) BLA to NAc (n = 11). Data are presented as mean ± SEM.

Finally, no significant differences were observed among groups in body weight or locomotor activity (Fig. 4A,B), indicating that the behavioral effects observed were not attributable to nonspecific alterations in general activity or physiological state.

**Figure 4.**
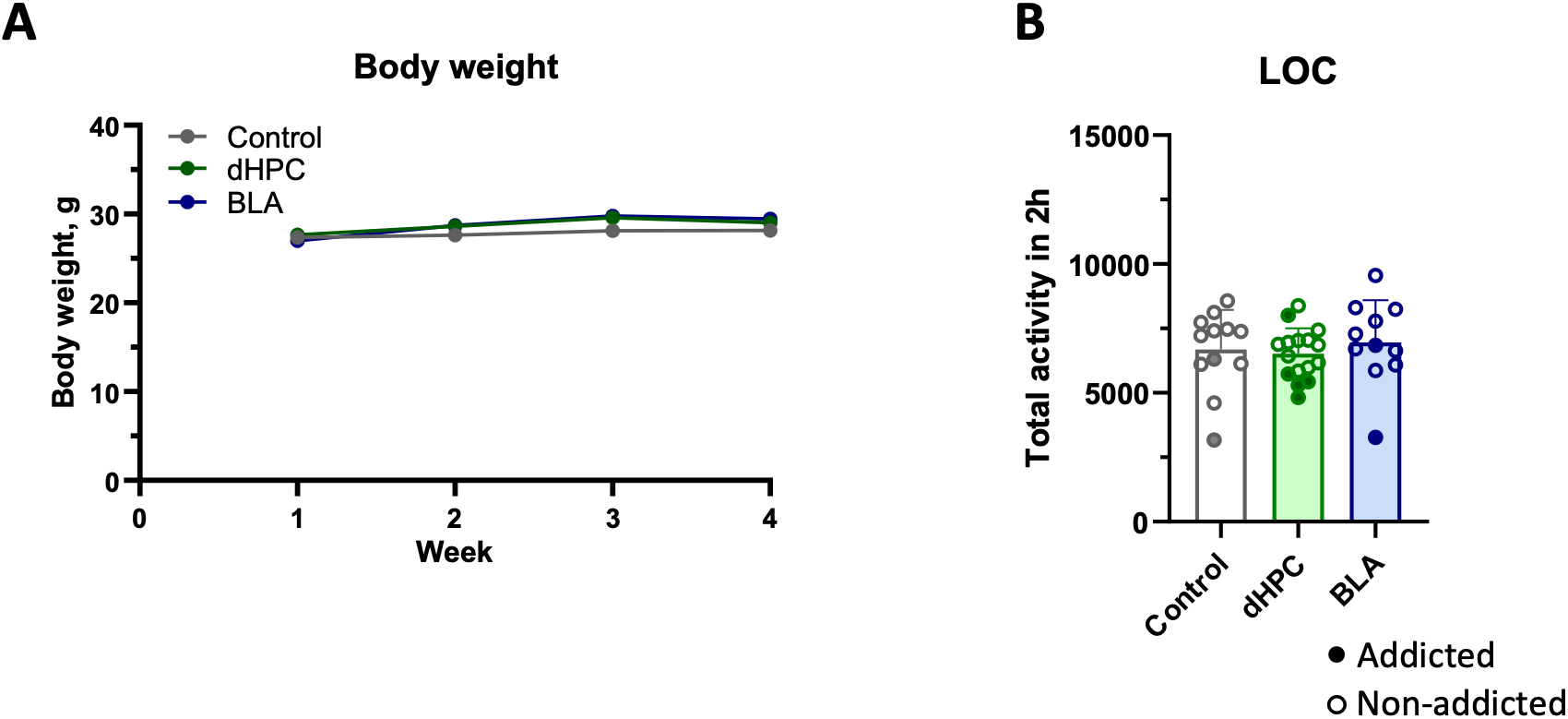
Chemogenetic inhibition of dHPC to NAc and BLA to NAc projections does not alter body weight or locomotor activity. **(A)** Body weight measured throughout the behavioral protocol. **(B)** Total locomotor activity during the 2-h locomotor activity test. Mice were randomly assigned to one of the three experimental groups: (1) Control (n = 12), (2) dHPC to NAc (n = 16), or (3) BLA to NAc (n = 11). Data are presented as mean ± SEM.

These findings suggest that dHPC to NAc inhibition preferentially enhances addiction vulnerability together with impulsivity, reward sensitivity, and resistance to extinction, whereas BLA to NAc inhibition mainly promotes persistence and cue-induced drug seeking.

## 4. DISCUSSION

The present study identifies two major glutamatergic afferents to the NAc, the dHPC to NAc and the BLA to NAc pathways, as key regulators of vulnerability to cannabinoid addiction-like behavior. Using a validated mouse model of WIN55,212-2 intravenous self-administration combined with projection-specific chemogenetic inhibition, we demonstrate that chronic suppression of either pathway markedly increased the proportion of mice developing an addiction-like phenotype. These findings reveal that glutamatergic inputs to the NAc exert a protective role against the transition from controlled cannabinoid use to addiction-like behavior.

Despite this common effect on addiction vulnerability, the two pathways differentially regulated specific behavioral components associated with the addiction process. Inhibition of both dHPC to NAc and BLA to NAc projections increased persistence of drug seeking during periods of drug unavailability, suggesting that these glutamatergic inputs converge on a shared neural mechanism that promotes behavioral control over drug seeking. In contrast, inhibition of the dHPC to NAc pathway selectively increased motivation to obtain the drug, impulsivity, reward sensitivity, and resistance to extinction, whereas inhibition of the BLA to NAc pathway preferentially enhanced cue-induced drug seeking. These findings indicate that distinct glutamatergic afferents to the NAc contribute to different behavioral dimensions underlying cannabinoid addiction vulnerability, supporting the view that the NAc integrates complementary contextual and emotional information to regulate the transition to compulsive drug use.

The NAc is a central interface between limbic, cortical, and motor systems, where motivationally relevant information is integrated to guide reward-directed behavior (Carlezon and Thomas, 2009). Rather than acting as a homogeneous structure, the NAc receives functionally specialized glutamatergic afferents from brain regions involved in contextual memory, emotional salience, executive control, and reward learning (Britt et al., 2012). These excitatory inputs provide the NAc with information about environmental context, conditioned cues, affective value, and action-outcome relationships, thereby shaping the selection and vigor of reward-seeking responses.

In the field of addiction, this integrative function becomes particularly relevant because drug-seeking behavior is driven not only by the pharmacological effects of the drug, but also by contextual and cue-associated information that acquires excessive motivational value (Kalivas and McFarland, 2003). In line with this view, our previous studies demonstrated that the PL to NAc pathway plays a critical role in compulsive-like behavior in food addiction (Domingo-Rodriguez et al., 2020), whereas inhibition of this same pathway did not significantly modify the main addiction-like criteria in the WIN55,212-2 model of cannabinoid addiction (Cajiao-Manrique et al., 2023b). These findings suggest that NAc-projecting glutamatergic circuits do not exert uniform effects across addictive disorders, but instead regulate specific behavioral dimensions depending on the reinforcer and the neurobiological context.

The present results extend this circuit-based framework by identifying two additional limbic glutamatergic inputs to the NAc, the dHPC to NAc and BLA to NAc pathways, as regulators of vulnerability to cannabinoid addiction-like behavior. Importantly, both pathways converged on persistence of drug seeking, while diverging in their contribution to motivation, impulsivity, reward sensitivity, extinction resistance, and cue-induced drug seeking. This suggests that the NAc integrates complementary contextual and emotional information through distinct glutamatergic pathways, and that disruption of these inputs facilitates the emergence of cannabinoid addiction-like behavior.

The dorsal hippocampus is traditionally associated with spatial and contextual memory, but growing evidence suggests it also contributes to reward-guided behavior by providing contextual information to the NAc (Volkow et al., 2019). Through its glutamatergic projections to the NAc, the dHPC encodes environmental contexts in which rewards are obtained, allowing reward-seeking behaviors to be appropriately adjusted according to the availability and predictability of reinforcement (Britt et al., 2012). Rather than simply promoting reward seeking, this pathway is thought to facilitate flexible, goal-directed behavior by integrating contextual information with motivational processes.

Our findings support this interpretation by showing that chronic inhibition of the dHPC to NAc pathway increased several behavioral dimensions associated with cannabinoid addiction vulnerability. In addition to increasing the proportion of mice classified as addicted, inhibition of this pathway enhanced persistence of drug seeking during periods of drug unavailability and increased motivation to obtain WIN55,212-2 under a progressive-ratio schedule. These findings suggest that intact dHPC to NAc signaling normally contributes to suppressing inappropriate drug-seeking behavior when reinforcement is unavailable, while limiting the excessive motivational value attributed to cannabinoid reward.

Interestingly, inhibition of the dHPC-to-NAc pathway also increased impulsivity and reward sensitivity, two behavioral traits associated with increased vulnerability to addiction. Elevated impulsivity has been proposed to facilitate the transition from controlled drug use to compulsive drug seeking, whereas enhanced reward sensitivity may increase the reinforcing efficacy of drugs of abuse (Belin et al., 2008). The concomitant increase in these phenotypic traits suggests that disruption of dHPC inputs to the NAc not only affects the expression of addiction-like behaviors but may also promote a behavioral profile that predisposes individuals to addiction.

Consistent with this interpretation, mice with inhibited dHPC to NAc projections also exhibited greater resistance to extinction, indicating a reduced ability to suppress previously acquired drug-seeking responses when the drug was no longer available (Bouton, 2004). Together with the increased persistence observed during the self-administration phase, these findings support the idea that the dHPC to NAc pathway plays a central role in updating reward-directed behavior according to changes in environmental contingencies. Loss of this contextual regulation may therefore contribute to the emergence of rigid and maladaptive drug-seeking behaviors that characterize the transition to cannabinoid addiction. Our findings suggest that the dHPC to NAc pathway is not promoting reward-seeking *per se*, but rather constraining it based on contextual information. In other words, the intact circuit appears to act as a brake, helping animals adjust their behavior when reinforcement contingencies change.

The BLA is a key component of the neural circuitry underlying emotional learning and the attribution of motivational value to environmental stimuli (Stefanik and Kalivas, 2013). Through its glutamatergic projections to the NAc, the BLA conveys information about the emotional significance of reward-associated cues, allowing conditioned stimuli to influence reward-seeking behavior (Ambroggi et al., 2008). This pathway has been consistently implicated in Pavlovian learning, cue-reward associations, and relapse to drug seeking triggered by conditioned cues across multiple drugs of abuse.

In the present study, chronic inhibition of the BLA to NAc pathway increased the proportion of mice classified as addicted and enhanced persistence of drug seeking during periods of drug unavailability, indicating that this circuit, similarly to the dHPC to NAc pathway, contributes to maintaining appropriate control over drug-seeking behavior. However, unlike dHPC inhibition, suppression of the BLA to NAc pathway did not alter motivation, impulsivity, reward sensitivity, or resistance to extinction. Instead, its most prominent behavioral effect was a selective increase in cue-induced drug seeking following extinction.

This dissociation is consistent with the established role of the BLA in assigning incentive salience to conditioned stimuli. Drug-associated cues progressively acquire motivational value through repeated pairings with drug reward, allowing them to trigger relapse even after prolonged periods of abstinence. Our findings suggest that intact glutamatergic signaling from the BLA to the NAc normally limits the capacity of conditioned cannabinoid-associated cues to regain control over behavior after extinction. Disruption of this pathway may therefore enhance the motivational impact of drug-paired cues, increasing the probability of relapse-like responding.

The observation that BLA to NAc inhibition selectively affected cue-induced drug seeking while leaving motivation under a progressive-ratio schedule unchanged further supports the notion that distinct neural mechanisms regulate the reinforcing properties of cannabinoids and the motivational influence of conditioned stimuli. Whereas motivation primarily reflects the willingness to work for drug reward, cue-induced reinstatement reflects the ability of previously learned environmental cues to reactivate drug seeking in the absence of drug availability. These findings therefore highlight a specific role of the BLA to NAc pathway in controlling cue-dependent aspects of cannabinoid addiction rather than reward valuation itself. Interestingly, the increase in cue-induced drug seeking following chronic BLA-to-NAc inhibition may appear to contrast with previous studies showing that acute inhibition of this pathway attenuates cue-induced cocaine seeking and conditioned reinforcement (Stefanik and Kalivas, 2013). However, the present manipulation was maintained throughout the development of addiction-like behavior rather than being restricted to the reinstatement test. Chronic suppression of the pathway may therefore have induced compensatory adaptations or disrupted the acquisition and updating of cue–outcome relationships, ultimately increasing cue control after extinction.

One of the most striking findings of the present study is that inhibition of both the dHPC to NAc and BLA to NAc pathways increased persistence of drug seeking during periods of drug unavailability, despite producing otherwise distinct behavioral phenotypes. Persistence is considered a core feature of addiction because it reflects the inability to suppress drug-seeking behavior when reward is unavailable, resembling the continued pursuit of drugs despite reduced or absent reinforcement observed in individuals with substance use disorders (Deroche-Gamonet et al., 2004). The fact that both pathways influenced this behavioral dimension suggests that persistence may represent a common output of NAc circuit function that integrates information from multiple glutamatergic afferents.

Although the dHPC and BLA convey different types of information to the NAc, contextual and emotional, respectively, both sources of information are required for the adaptive regulation of reward-seeking behavior. Under physiological conditions, contextual information allows animals to determine whether drug seeking is appropriate in a given environment, whereas emotionally salient cues help predict reward availability based on previous experience. Integration of these complementary signals within the NAc enables flexible behavioral responses that are continuously updated according to environmental contingencies. Disruption of either input may therefore impair this integrative process, leading to persistent drug seeking even when reinforcement is unavailable.

Interestingly, persistence was the only addiction-like criterion altered by inhibition of both pathways, whereas motivation, impulsivity, reward sensitivity, extinction resistance, and cue-induced drug seeking showed pathway-specific regulation. This dissociation suggests that persistence may constitute a central behavioral process through which multiple limbic circuits influence vulnerability to cannabinoid addiction, while other addiction-related behaviors are controlled by more specialized neural networks. These findings support a model in which the NAc functions as a hub that integrates complementary glutamatergic information to regulate distinct components of addictive behavior, with persistence emerging as a shared circuit-level output of this integration. Cannabis use disorder (CUD) is increasingly recognized as a major public health concern, yet its underlying circuit-level mechanisms remain poorly understood and no pharmacological treatment has demonstrated consistent clinical efficacy (Spiga et al., 2025). A major obstacle to the development of effective therapies is the limited understanding of the neural substrates that confer vulnerability to the transition from recreational cannabis use to compulsive drug seeking. By identifying specific glutamatergic afferents to the NAc that regulate distinct behavioral dimensions of cannabinoid addiction, the present study provides new insight into the neural circuitry underlying this transition.

Our findings suggest that cannabinoid addiction does not arise from dysfunction of a single brain region, but rather from alterations in distributed limbic circuits that converge on the NAc. Importantly, the observation that the dHPC to NAc and BLA to NAc pathways regulate complementary aspects of addiction-like behavior indicates that different symptoms of CUD may depend on partially dissociable neural substrates. Whereas disruption of the dHPC to NAc pathway primarily affected motivational and contextual control over drug seeking, inhibition of the BLA to NAc pathway preferentially enhanced the influence of conditioned drug-associated cues. This functional dissociation may help explain the heterogeneity observed among individuals with CUD, in whom craving, cue reactivity, compulsive use, and relapse vulnerability often present with different degrees of severity.

These findings also have potential therapeutic implications. Current interventions for CUD are largely symptom-oriented and do not specifically target the neural circuits that drive maladaptive drug-seeking behavior. Although direct manipulation of these pathways is not currently feasible in clinical practice, identifying circuit-specific mechanisms provides a framework for the development of more selective therapeutic strategies. Future approaches aimed at restoring the activity of specific glutamatergic projections to the NAc, either through pharmacological modulation, neuromodulation techniques, or circuit-informed behavioral interventions, may offer new opportunities to reduce vulnerability to cannabinoid addiction and prevent relapse.

More broadly, the present work supports the concept that addiction should be understood as a disorder of dysfunctional neural circuits rather than isolated brain regions. Identifying how individual glutamatergic pathways contribute to specific behavioral dimensions of addiction represents an important step toward developing mechanism-based interventions capable of targeting the distinct processes that underlie Cannabis Use Disorder.

## Acknowledgements

This work was supported by Spanish “Ministerio de Ciencia, Innovación y Universidades, Agencia Estatal de Investigación (AEI)” (PID2020-120029GB-I00/MICIN/AEI/10.13039/501100011033, RD21/0009/0019 and PDI2023-1511680B-C21), the Spanish “Instituto de Salud Carlos III, RETICS-RTA” (#RD12/0028/0023), the “Generalitat de Catalunya, AGAUR” (#2020 SGR), “ICREA-Acadèmia” (#2025) and the Spanish “Ministerio de Sanidad, Servicios Sociales e Igualdad”, “Plan Nacional Sobre Drogas of the Spanish Ministry of Health” (#PNSD-2022) to R.M., “la Caixa Health” LCR/PR/HR22/5240017 to R.M. and E.M-G., “Plan Nacional Sobre Drogas of the Spanish Ministry of Health” (#PNSD-2019I006, #PNSD-2023I040) and Spanish “Ministerio de Ciencia e Innovación” (ERA-NET) PCI2021-122073-2A to E.M-G. We are very grateful to R. Martín and F. Porrón for their technical support. Figure 1 was created with BioRender.

## Author contributions

E.M.-G. and R.M. conceived and designed the behavioral studies. T.G., M.F.P.-B., and R.C. performed the behavioral experiments and statistical analyses under the supervision of E.M.-G. and R.M. The intravenous catheterization surgeries were performed by E.M.-G. with assistance from T.G., M.F.P.-B., and R.C. R.C. performed brain dissection. I.M.B performed the histological validation of viral injections. M.G.F. and R.A. performed RNA extraction. Next, T.G., M.F.P.-B., R.C., E.M.-G., and R.M. wrote the manuscript. Finally, R.M., E.M.-G., and I.M.B. critically revised the manuscript with input from all authors.

## Data availability

Individual data points are graphed in the main figures. All the relevant data that support this study are available from the corresponding author to any interested researcher upon reasonable request.

